# Pre-stimulus pupil-linked arousal enhances initial stimulus availability and accelerates decay in iconic memory

**DOI:** 10.1101/2025.10.31.685757

**Authors:** Paul Justin Connor Smith, Niko A. Busch

## Abstract

Pupil diameter reflects not only reflexive responses to light but also spontaneous fluctuations in arousal. Such pre-stimulus fluctuations may shape visual performance, yet it remains unclear whether they only affect perceptual sensitivity or decision processes in near-threshold tasks, or instead influence the temporal availability of sensory information even for highly visible stimuli. We tested whether pupil-linked arousal modulates iconic memory, a high-capacity but short-lived visual store bridging sensory encoding and short-term memory. Based on a model in which a brief stimulus evokes an internal response decaying over a few hundred milliseconds, we hypothesized that increased arousal enhances performance either by amplifying the initial response or by slowing its decay. Thirty-seven participants viewed brief displays of six oriented stimuli, followed by a cue after a variable stimulus–cue onset asynchrony indicating which item to report. Pre-stimulus pupil diameter was continuously recorded, and behavioral data were fitted with an exponential decay function estimating initial stimulus availability, decay rate, and asymptotic performance. Larger pre-stimulus pupil dilation was associated with higher initial stimulus availability but faster decay of the iconic trace, while information transferred into short-term memory and confidence ratings remained unaffected. These findings demonstrate that spontaneous fluctuations in arousal shape the temporal dynamics of sensory persistence. Elevated arousal boosts the initial strength of visual representations but shortens their duration, suggesting that arousal adaptively prioritizes rapid updating of visual input over prolonged stability.

**Significance Statement:** How does arousal shape the earliest stages of visual memory? By leveraging a criterion-free partial-report task that emphasizes the temporal availability of sensory information, we show that spontaneous pre-stimulus pupil dilation – an index of arousal – selectively boosts the initial availability of iconic memory while accelerating its decay. This dual effect suggests that arousal tunes the temporal dynamics of sensory persistence, potentially prioritizing rapid updating of incoming information. These findings advance the understanding of how state-dependent fluctuations in arousal shape early perception in a context that minimizes decision-bias confounds.

## Introduction

Pupil size is influenced by both external and internal factors. Constriction in response to increased brightness or reduced viewing distance serves to optimize visual acuity (Mathôt & Ivanov, 2019). Beyond these reflexive adjustments, pupil dilation reflects internal states, such as heightened emotional, physical, or cognitive arousal (Hess & Polt, 1960, 1964; Kahneman, 1973; Mathôt, 2018; Stanners et al., 1979). This arousal-related dilation arises from phasic norepinephrine release by the locus coeruleus (Aston-Jones & Cohen, 2005; Joshi & Gold, 2020; Joshi et al., 2016), which modulates a range of cognitive processes, including attention, memory, and vision (Bradley et al., 2008; Kahneman & Beatty, 1966; Laeng et al., 2012; Robbins, 2000; Stanners et al., 1979). The resulting relationship between arousal and behavioral performance typically follows an inverted U-shape, with optimal performance benefits emerging at intermediate arousal levels (Aston-Jones & Cohen, 2005). Within this range, higher arousal is associated with better visual task performance and larger pupils (Eberhardt et al., 2022; Mathôt, 2020; Pilipenko & Samaha, 2024), likely reflecting enhanced visual sensitivity and attentional allocation (Alnæs et al., 2014; Irons et al., 2017; Mathôt, 2018; Mathôt & Ivanov, 2019). The dominant theory posits that increases in arousal broaden the focus of attention, thereby improving both performance and metacognitive measures in visual tasks (Hoeks & Ellenbroek, 1993; Irons et al., 2017; Kolnes et al., 2024). Previous research has primarily examined pupil responses following stimulus presentation, but recent work has begun to explore the influence of *pre-stimulus* pupil fluctuations on visual processing (Grujic et al., 2024). Most of these studies have focused on detection tasks, in which performance is limited by the visibility of stimulus presence (Eberhardt et al., 2022; Pilipenko & Samaha, 2024), or discrimination tasks, where performance is limited by the discriminability of stimulus features (Allen et al., 2016). Detection paradigms often yield ambiguous results because changes in detection rates can reflect genuinely improved perceptual sensitivity (i.e., better discrimination between signal and noise), strategic shifts in decision criterion, or changes in subjective signal-likeness (Iemi et al., 2017; Samaha et al., 2020). Moreover, it remains unclear whether pupil-linked arousal affects performance only in tasks using near-threshold stimuli—where performance is constrained by low visibility or discriminability—or whether its influence extends to tasks in which performance is instead limited by the persistence of stimulus information.

To address this question, we tested the effect of pre-stimulus pupil size on performance in an iconic memory task—a high-capacity but short-lived visual memory store that depends on persistent neuronal activity in early visual cortex (Teeuwen et al., 2021). Iconic memory represents the immediate sensory trace that follows visual stimulation and precedes visual short-term memory, which is more durable but has much lower capacity (Dick, 1974; Sperling, 1960). On each trial, a display of six concentric stimuli was briefly flashed, followed by a cue indicating which stimulus to report after a variable stimulus–cue onset asynchrony (SOA) (Lu et al., 2005). Participants reported the orientation of the cued stimulus.

Importantly, stimuli in the iconic memory partial-report task are clearly visible, and their orientations are easily discriminable. Performance in this task is limited not by visibility but by the capacity of short-term memory, which can hold only a small number of items. Accordingly, when the report cue appears simultaneously with the display, participants can immediately select a single stimulus and maintain it in short-term memory, leading to near-perfect performance. As the SOA increases, performance declines because selection and transfer into short-term memory must rely on the rapidly fading iconic trace. This trace persists for only a few hundred milliseconds, after which detailed visual information is no longer accessible. The non-linear drop in performance at intermediate SOAs thus reflects the short-lived availability of detailed stimulus information once the display has disappeared. Consequently, the psychometric decay function describing performance across SOAs provides a quantitative measure of the persistence of visual information (Gegenfurtner & Sperling, 1993) and a sensitive tool for studying how pupil-linked arousal modulates temporal stimulus availability.

We hypothesized that increases in spontaneous pre-stimulus pupil dilation, indicating fluctuations in arousal, result in increased performance in the partial-report paradigm. Specifically, iconic memory performance can be modulated by two main factors: the strength of the initial stimulus representation and the speed of the decay of the memory information (Figure 1B). Our study allows us to test whether pupil-linked arousal impacts this initial stimulus availability or the decay of the memory trace. This would further cement the link between arousal and visual performance, in a paradigm which minimizes decision-bias confounds, whilst also establishing a novel link between spontaneous arousal and its specific impact on iconic memory performance.

**Figure 1:**
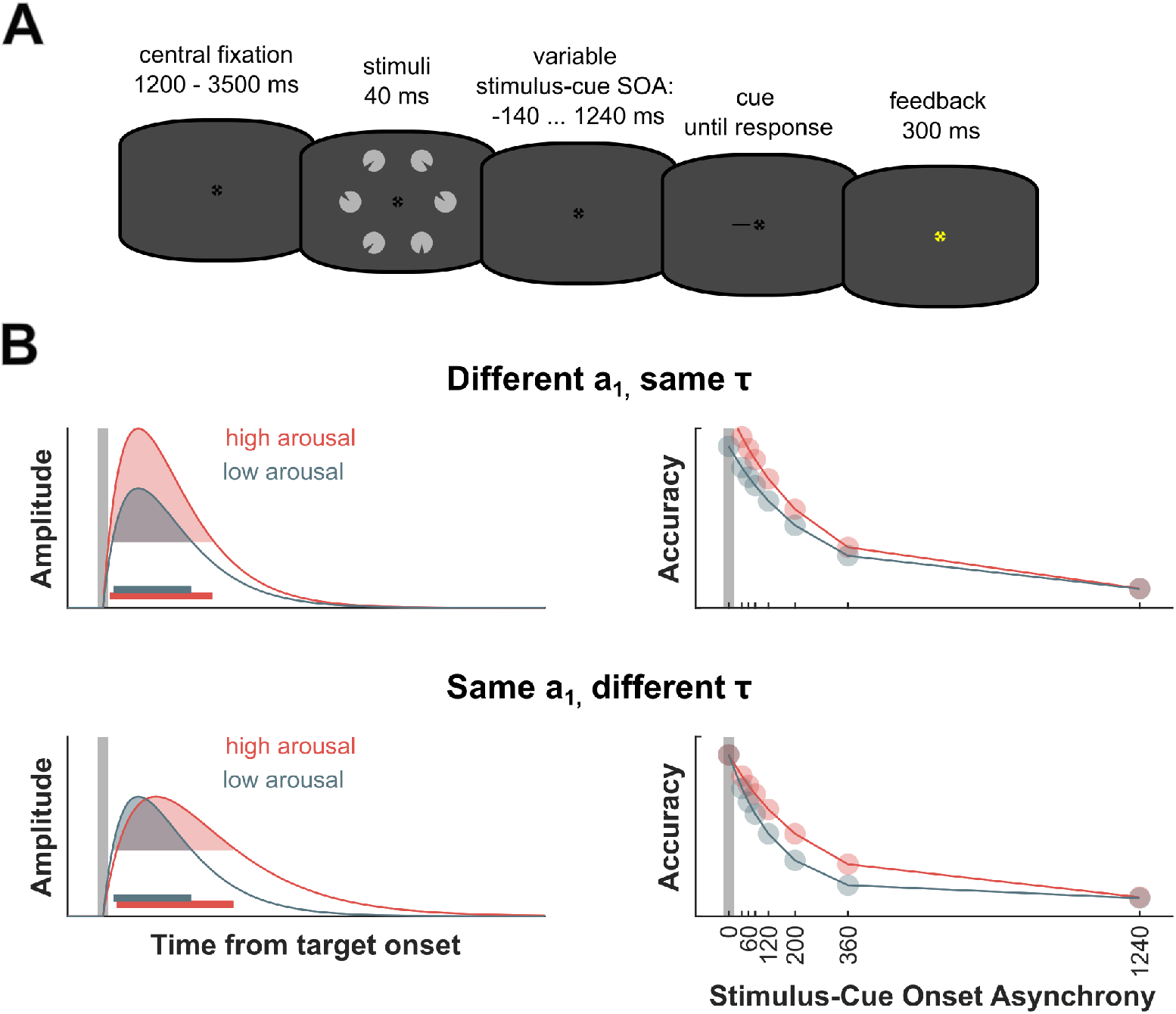
**A**: Illustration of the partial report paradigm. Each trial began with a central fixation cross displayed for a variable interval of 1200-3500 ms. Following fixation, a stimulus array consisting of six items was presented for 40 ms. Participants were then cued to report the orientation of one target item, with the cue appearing at various SOAs relative to stimulus onset: 140 ms before stimulus onset, at stimulus onset, or 40, 60, 80, 120, 200, 360, or 1240 ms after. There was no time limit for reporting or the subsequent confidence rating. After the confidence rating, feedback was given via the color of the fixation cross, turning blue for a correct response and yellow for an incorrect one. **B**: Illustration of two possible mechanisms of temporal persistence modulation. Left: Schematic representation of the impulse response triggered by stimulus onset (indicated by the vertical gray bar). The response shows a rapid onset followed by a gradual decay. The stimulus information persists as long as the response amplitude remains above a critical threshold (shaded area). Horizontal bars indicate duration of persistence. Heightened arousal can extend this persistence either by increasing the *amplitude* of the initial response (top) or by slowing its *decay* (bottom). Right: Hypothetical performance in the partial report task as a function of stimulus-cue SOA. Performance is high at short SOAs and declines with longer SOAs. An amplified response (top) is predicted to enhance initial stimulus availability, thereby boosting performance primarily at short SOAs, which would be captured by the model’s *a*_1_ parameter (see equation 1). A slower decay (bottom) would improve performance at intermediate SOAs, reflected in the model’s *τ* parameter.

We hypothesized that spontaneous fluctuations in pre-stimulus pupil dilation – reflecting changes in arousal – modulate performance in the partial-report paradigm. Specifically, we assume that a brief visual stimulus elicits an internal impulse response that outlasts the external stimulus due to the visual system’s low-pass properties (Di Lollo, 1980; Loftus & Irwin, 1998). Stimulus information remains available for report as long as this response, corresponding to the iconic trace, stays above a critical threshold. Increased arousal could extend this period of availability in two distinct ways (Figure 1B): either by amplifying the initial response, thereby increasing the strength of the initial stimulus representation, or by slowing its decay, thereby prolonging the persistence of the iconic trace. These mechanisms would produce dissociable patterns in the partial-report task: amplitude changes mainly affecting performance at short stimulus-cue SOAs, and decay changes affecting performance at intermediate delays. This approach strengthens the link between arousal and perceptual performance in a paradigm that minimizes decision-bias confounds and directly quantifies temporal stimulus availability.

## Material and Methods

### Pre-registration

The study was pre-registered as part of a larger project on the Open Science Framework (osf.io/u6h8d). The analyses of pupil data reported here were conducted exploratory and were not part of the original preregistered plan.

### Participants

EEG and eye-tracking data were collected from 61 healthy participants (mean age = 23.6 ± 3.9 years; 45 female). The final sample comprised 37 participants (mean age = 23.1 ± 3.3 years; 26 female) after excluding five for poor performance (< 80% accuracy at −140 ms and 0 ms SOA), 14 for incomplete eye-tracking data, and eight whose data could not be fit by the exponential decay model (not preregistered; see below). All participants had normal or corrected-to-normal vision, reported no neurological or psychiatric disorders, provided written informed consent, and received course credit or monetary compensation. The study was approved by the ethics committee of the University of Münster (Ref. 2023-27-PS).

### Stimuli and Procedure

The experiment was programmed in MATLAB 2022 (MathWorks) using Psychtoolbox (Brainard & Vision, 1997; Kleiner et al., 2007; Pelli, 1997) and presented on a 24” Viewpixx/EEG LCD monitor (1920×1080 px, 120 Hz, 1 ms response time, 95% luminance uniformity; www.vpixx.com). Background luminance during the pre-stimulus interval was 45.24 cd/m^2^, and the fixation cross had a luminance of 0.16 cd/m^2^.

Each trial in this partial-report paradigm began with a central fixation cross (0.6°, RGB [0 0 0]) shown for 1200–3500 ms, followed by six stimuli arranged in a circle and presented for 40 ms. Stimuli were light-grey circles (2.6° visual angle, RGB [180 180 180]) with a wedge (0.28° thick, 0.6° diameter) cut out at one of eight orientations (0°–315° in 45° steps). A black line (1° length, 0.1° thickness) served as the post-cue, pointing to one of the stimulus locations. The cue appeared after a variable SOA of either 140 ms before stimuli onset, at stimuli onset, or 40, 60, 80, 120, 200, 360, or 1240 ms after stimuli onset. Participants were instructed to press a number on the number pad corresponding to the orientation of the wedge in the target circle (eight for 0°, six for 90°, two for 180°, etc.). Following this decision, the participants were instructed to press either four (low), five (medium), or six (high) on the number pad, indicating their respective confidence rating. There was no time limit for their response. Feedback on their decision was provided 300 ms after their confidence rating, with correct responses marked by a blue (RGB: [0 0 255]) fixation cross, and incorrect responses indicated by a yellow (RGB: [255 255 0]) fixation cross. Participants were instructed to avoid eye movements and blinks during the stimulus presentation and to fixate on the center of the screen. A visualization of a trial can be seen in Figure 1A. There were 1296 trials, with short self-paced breaks after 170 consecutive trials, with counterbalanced SOA, stimulus position around the fixation cross, and target cut-out orientation.

### Acquisition and Analysis of Pupil Data

Recordings took place in a dimly lit, soundproof cabin (illuminance during the pre-stimulus interval: 17 lux). Participants’ heads were stabilized on a chinrest at a viewing distance of approximately 86 cm. Eye movements were monitored with a desktop-mounted EyeLink 1000+ infrared eye tracker (SR Research Ltd.) sampling at 1000 Hz (monocular, dominant eye). Data were recorded in.edf format, and pupil dilation was measured in the default EyeLink unit (pixels).

All eye-tracking data preprocessing and analysis steps were conducted in MATLAB 2023a (mathworks.com) using the EEGLAB toolbox (Delorme & Makeig, 2004) and custom scripts. Trials containing eye blinks within a −500 to +500 ms window around stimulus onset, or instances where gaze deviated more than 2.5° of visual angle (dva) from the fixation cross, were rejected. On average, 29.3 trials were excluded per participant.

Pre-stimulus pupil dilation was computed for each trial as the average pupil size within a window from −500 to −2 ms before stimulus onset. A subtractive baseline correction was applied using a baseline interval from −700 to −501 ms. Baseline-corrected pupil data were z-scored within subjects, and trials with pupil values exceeding *±* standard deviations were excluded from further analysis (Mathôt & Vilotijević, 2023; Mathôt et al., 2018). On average, 53.3 trials were removed per participant. Singletrial pupil dilation values were then divided into three quantile-based bins to examine a potential inverse U-shaped relationship between arousal and performance. Performance and confidence data were averaged within each pupil bin for all non-negative stimulus–cue SOAs.

### Modeling the Time Course of Iconic Memory

This study aimed to test the hypothesis that pre-stimulus arousal modulates the temporal availability of stimulus information in iconic memory. We assumed that a brief visual stimulus evokes an impulse response that outlasts the physical stimulus due to the low-pass filtering properties of the early visual system (Di Lollo, 1980; Loftus & Irwin, 1998). The longer this internal response remains above a critical threshold (shaded areas in Figure 1B), the longer stimulus information persists and remains available for perceptual and decisional processes. Greater arousal may enhance this persistence either by amplifying the initial response amplitude (Figure 1B, top) or by slowing its decay (Figure 1B, bottom).

To test how pupil-linked arousal affects these two aspects of stimulus persistence, we analyzed behavioral sensitivity (*d*′) and confidence across stimulus–cue SOAs. Confidence values were first normalized to a 0–1 scale. For each participant, both measures were fitted with a non-linear exponential decay function,

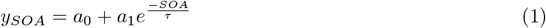

where *y* denotes either *d*′ or confidence, using a nonlinear least-squares solver with starting parameters based on Lu et al. (Lu et al., 2005). The model included three parameters: *a*_0_, representing sensitivity at long SOAs (information transferred into short-term memory without cueing benefits); *a*_1_, representing the initial, fast-decaying availability of stimulus information; and *τ*, the time constant of this decay. Model fitting was first applied to trials from all non-negative stimulus–cue SOAs.

Model fitting was first applied to all trials prior to binning by pupil dilation, using unconstrained model parameters. Participants with parameter estimates exceeding three standard deviations from the group mean in either measure were excluded from further analyses of that measure (see Participants). Subsequently, the model was fitted separately to *d*′ and confidence data separately for each pupil-dilation bin, using the best-fitting parameters from the previous step as starting values. The fitted confidence functions for the large pre-stimulus pupil dilation bin contained several extreme outliers, with parameter values several orders of magnitude larger than the group average. Because these outliers were not identified in the overall confidence fits (see Participants), three participants with parameter values exceeding 10 standard deviations from the group mean were removed from the confidence modeling. This reduced the sample for the confidence analysis to 34 participants (mean age = 22.91 ± 3.34 years; 24 female, 10 male). Although we suspect that additional outliers may still influence the fit, we decided against arbitrarily removing participants who did not exhibit parameter values larger than 10 times the average parameter value.

### Statistical Analysis

Deviating from the preregistration, we chose a simplified analysis approach in place of the originally planned jackknife method. To test the relationship between pre-stimulus pupil dilation and iconic memory performance, the model parameters *a*_0_, *a*_1_, and *τ* were analyzed using one-way repeated-measures ANOVAs with the within-subject factor pupil dilation bin (three levels: low, medium, high). The same analysis was applied to the confidence data. Pairwise post-hoc two-tailed t-tests were used to compare individual bins.

### Data and code accessibility

The data will be available upon acceptance of the manuscript at the Open Science Framework (osf.io/ xfkrm/). The code will be available at github.com/pauljcs/alpha-iconic.

## Results

### Performance Data

The time-course of pupil size in the pre-stimulus time window (−500 to −2 ms before stimulus onset) indicated that trials with correct responses were preceded by larger pupil size. This effect began to emerge approximately 300 ms before stimulus onset (see Figure 4). Initial stimulus availability (*a*_1_) differed significantly between pupil dilation bins, (*F* (2, 72) = 4.07, *p* = 0.012, (see Figure 2A, Figure 3A, and Table 1). Post-hoc t-tests showed that *a*_1_ was significantly higher for large vs. small pre-stimulus pupil dilation (*p* = 0.046) and for medium vs. small pre-stimulus pupil dilation (*p* = 0.002), with no other significant differences between bins (see Table 2).

**Table 1:** *d*′ Function Parameters ANOVA.

| Parameter | Effect | SumSq | $df$ | MeanSq | $F$ | $p$ | $\eta_p^2$ |
| --- | --- | --- | --- | --- | --- | --- | --- |
| $a_0$ | Bins | 0.2589 | 2 | 0.1294 | 1.57 | .215 | .041 |
|  | Error | 5.9412 | 72 | 0.0825 |  |  |  |
| $a_1$ | Bins | 1.7575 | 2 | 0.8788 | 4.70 | .012* | .115 |
|  | Error | 13.459 | 72 | 0.1869 |  |  |  |
| $\tau$ | Bins | 0.2002 | 2 | 0.1001 | 5.14 | .008** | .125 |
|  | Error | 1.4032 | 72 | 0.0195 |  |  |  |

Repeated-measures ANOVA results for the exponential-decay function parameters across three pre-stimulus pupil-dilation bins (small, medium, large) for the $d'$ . The first column lists the parameter ( $a_0$ , $a_1$ , $\tau$ ). The second column indicates the effect (Bins, Error). The remaining columns report the sum of squares (SumSq), degrees of freedom ( $df$ ), mean square (MeanSq), the $F$ -statistic, the $p$ -value, and partial eta squared ( $\eta_p^2$ ). Significant effects are marked with stars (\* = $p < 0.05$ ; \*\* = $p < 0.01$ ).

**Table 2:** *d*′ Function Parameters Post-hoc Pairwise Comparisons (Mean Differences)

| Parameter | Comparison | Difference | $SE$ | $p$ | 95% CI Lower | 95% CI Upper |
| --- | --- | --- | --- | --- | --- | --- |
| $a_0$ | small - medium | -0.084 | 0.067 | .218 | -0.220 | 0.052 |
|  | small - large | -0.114 | 0.066 | .092 | -0.248 | 0.019 |
|  | medium - large | 0.030 | 0.068 | .656 | -0.167 | 0.107 |
| $a_1$ | small - medium | -0.289 | 0.089 | .003** | -0.470 | -0.108 |
|  | small - large | -0.237 | 0.115 | .046* | -0.470 | -0.004 |
|  | medium - large | 0.052 | 0.096 | .592 | -0.143 | 0.247 |
| $\tau$ | small - medium | 0.068 | 0.038 | .084 | -0.010 | 0.146 |
|  | small - large | 0.102 | 0.032 | .003** | 0.037 | 0.167 |
|  | medium - large | -0.034 | 0.026 | .193 | -0.086 | 0.086 |

Post-hoc pairwise comparison t-test results for the exponential-decay function parameters of $d'$ across pre-stimulus pupil-dilation bins. The first column lists the parameter ( $a_0$ , $a_1$ , $\tau$ ). The second column indicates the bin contrast (small-medium, small-large, medium-large). The third and fourth columns report the mean difference between bins and its standard error (SE). The fifth column shows the $p$ -value. The sixth and seventh columns provide the lower and upper bounds of the 95% confidence interval for the difference. Negative differences indicate lower values in the first bin relative to the second. Significant comparisons are marked with stars (\* = $p < 0.05$ ; \*\* = $p < 0.01$ ).

**Figure 2:**
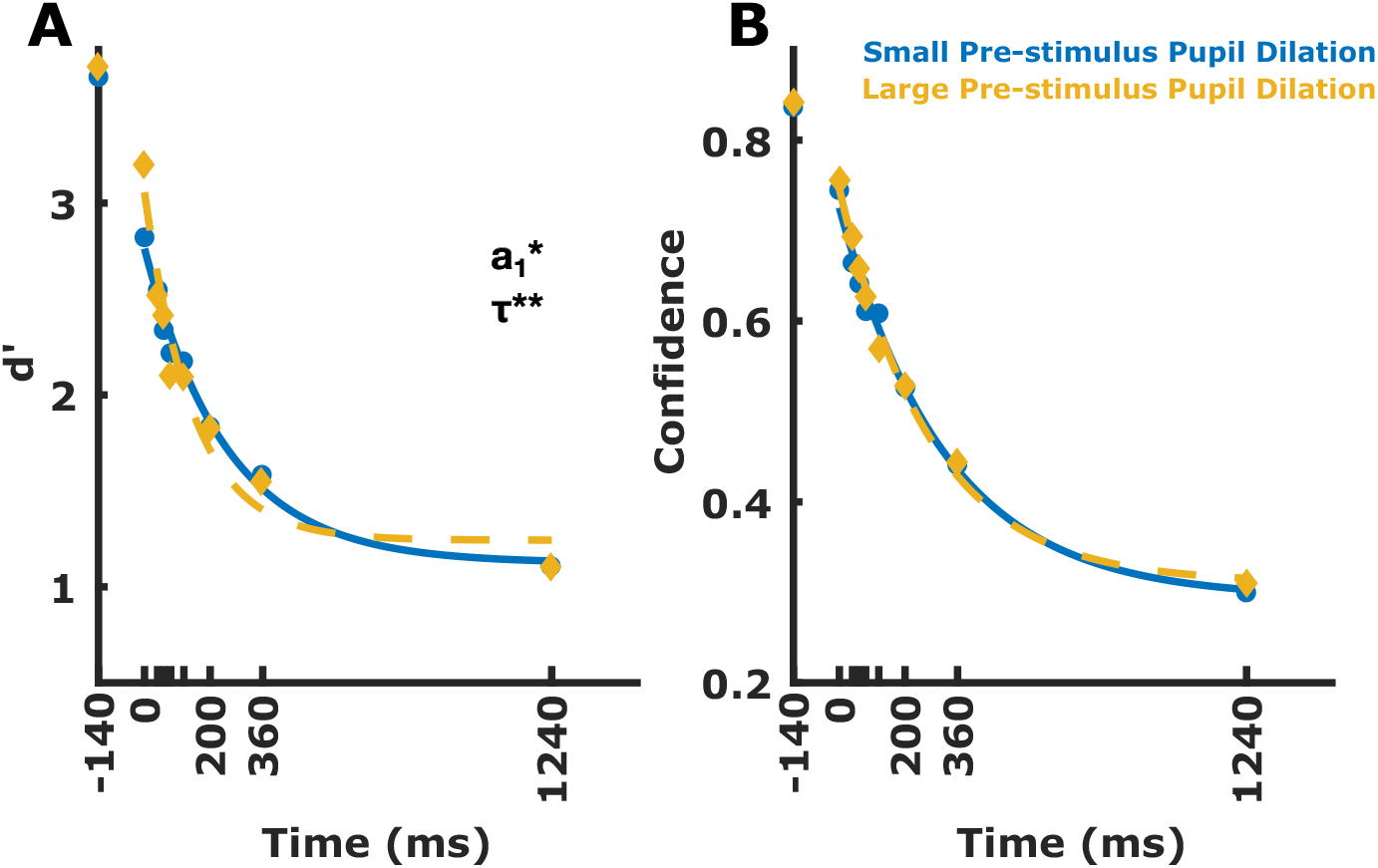
Accuracy (*d*′) and normalized confidence data for each SOA. Lines show model fits. Data are shown separately for **A**: accuracy for trials with large vs. small pre-stimulus pupil dilation & **B**: confidence for trials with large vs. small pre-stimulus pupil dilation. The blue line indicates the small pre-stimulus pupil dilation data, the red line indicates the medium pre-stimulus pupil dilation data, and the yellow line indicates the large pre-stimulus pupil dilation data. Model parameters showing a significant difference between the conditions are indicated with an asterisk. For illustration purposes, the medium pupil dilation bin is not shown.

**Figure 3:**
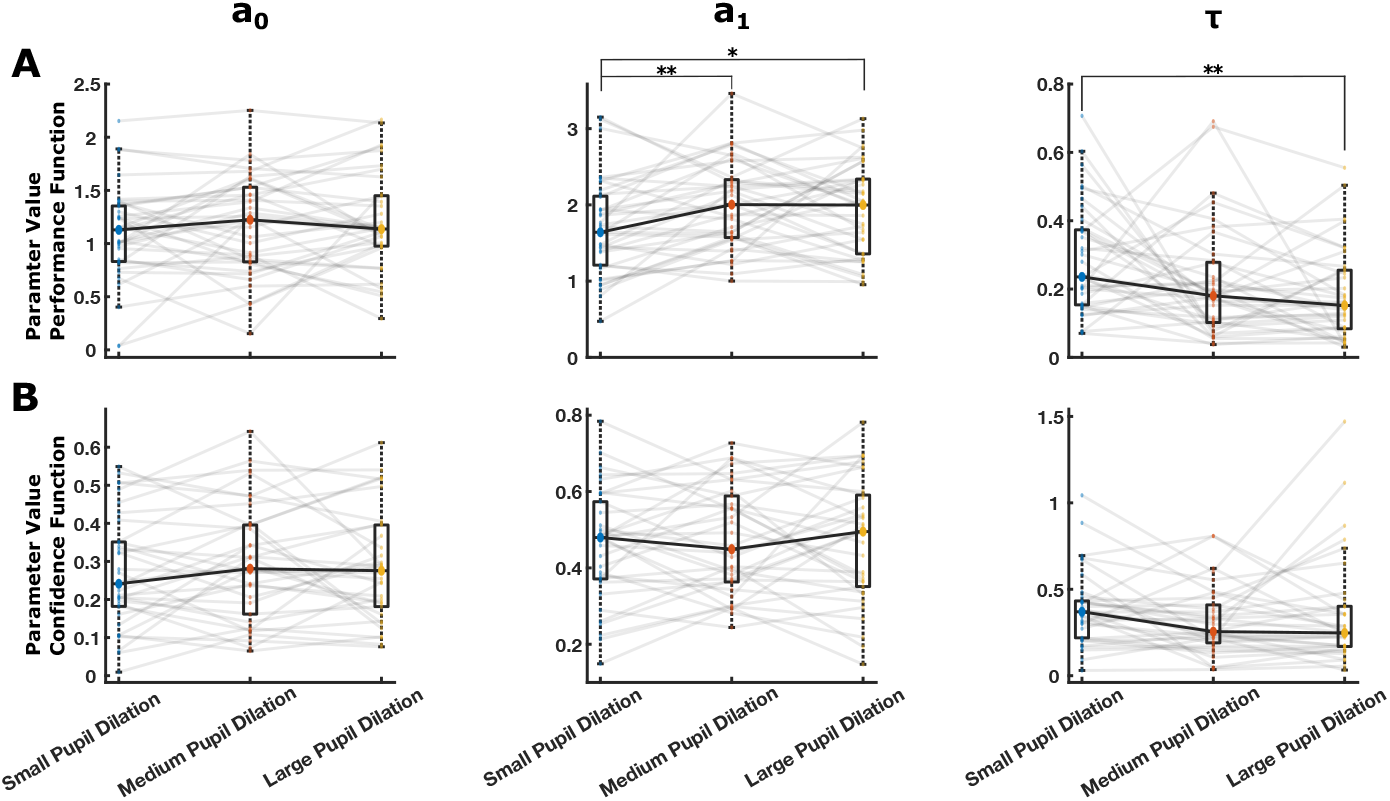
Function parameter (a_0_, a_1_ and *τ*) differences between small, medium and large pre-stimulus pupil dilation bins for **A**: accuracy (*d*′) data and **B**: normalised confidence data. Boxplots for small, medium and large pre-stimulus pupil dilation are shown. The large blue dot represents the median parameter value for small pre-stimulus pupil dilation, the large red dot represents the median parameter value for small pre-stimulus pupil dilation, and the large yellow dot represents the median parameter value for large pre-stimulus pupil dilation. Small dots represent the individual participants’ function parameter values. Solid lines show the median parameter value difference. Transparent lines show the parameter value difference for individual participants. Model parameters showing a significant difference between the conditions are indicated with an asterisk (* = *p <* 0.05; ** = *p <* 0.01).

**Figure 4:**
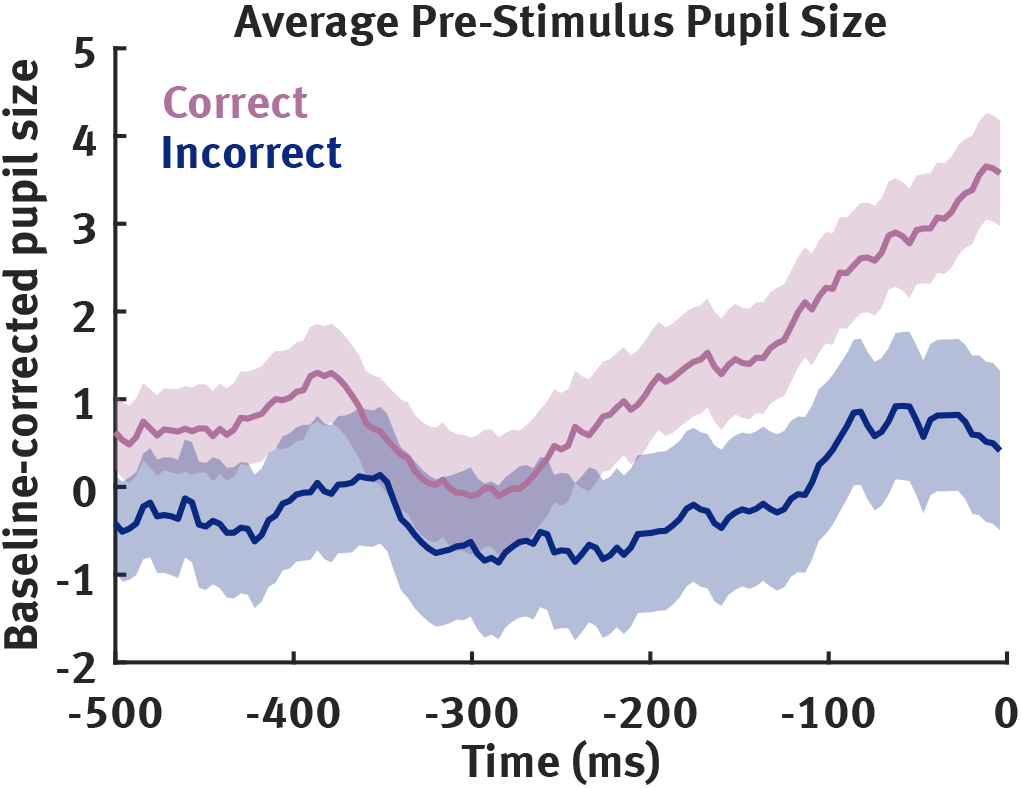
Average baseline corrected pupil size (arbitrary EyeLink unit) in the pre-stimulus time window (−500 to −2 ms before stimulus onset). The purple line indicates the pupil size for correct responses, and the dark blue line indicates the pupil size for incorrect responses. Shaded areas represent the SEM.

The time constant of iconic decay (*τ*) also differed significantly between pupil dilation bins, *F* (2, 72) = 5.136, *p* = 0.008 (see Figure 2A, Figure 3A, and Table 1). Post-hoc t-tests revealed that *τ* was significantly smaller for large than for small pre-stimulus pupil dilation (*p* = 0.003) with no other significant differences between bins (see Table 2).

The asymptotic parameter (*a*_0_), reflecting information transferred to short-term memory, showed no significant effect of pupil dilation bin, *F* (2, 72) = 1.569, *p* = 0.215 (see Figure 2A, Figure 3A, and Table 1).

### Confidence Data

The function parameters representing initial stimulus availability (*a*_1_), information transfer to short-term memory (*a*_0_), and the speed of iconic decay (*τ*) did not differ significantly between pupil dilation bins (*a*_1_: *F* (2, 66) = 0.20, *p* =.851; *a*_0_: *F* (2, 66) = 0.74, *p* =.481; *τ* : *F* (2, 66) = 1.55, *p* =.220; see Figures 2B and 3B; Table 3).

**Table 3:** Confidence Parameters ANOVA.

| Parameter | Effect | SumSq | $df$ | MeanSq | $F$ | $p$ | $\eta_p^2$ |
| --- | --- | --- | --- | --- | --- | --- | --- |
| $a_0$ | Bins | 0.0097 | 2 | 0.00487 | 0.74 | .481 | .022 |
|  | Error | 0.4337 | 66 | 0.00657 |  |  |  |
| $a_1$ | Bins | 0.0036 | 2 | 0.00180 | 0.20 | .815 | .006 |
|  | Error | 0.5802 | 66 | 0.00879 |  |  |  |
| $\tau$ | Bins | 0.0908 | 2 | 0.04541 | 1.55 | .220 | .045 |
|  | Error | 1.937 | 66 | 0.02935 |  |  |  |

Repeated-measures ANOVA results for the exponential-decay confidence-function parameters across three pre-stimulus pupil-dilation bins (low, medium, high). The first column lists the parameter ( $a_0$ , $a_1$ , $\tau$ ). The second column indicates the effect (Bins, Error). The remaining columns report the sum of squares (SumSq), degrees of freedom ( $df$ ), mean square (MeanSq), the $F$ -statistic, the $p$ -value, and partial eta squared ( $\eta_p^2$ ).

## Discussion

Do spontaneous changes in pupil-linked arousal influence iconic memory performance? Our study addressed this question by examining pre-stimulus fluctuations in pupil dilation and relating these variations to performance in a partial-report paradigm. We hypothesized that larger pre-stimulus pupil dilation, reflecting heightened physiological arousal, would enhance iconic memory performance by modulating either the initial availability of stimulus information or the rate of its decay (Figure 1B). This hypothesis builds on previous work showing that increased pupil dilation predicts better performance in visual detection tasks (Eberhardt et al., 2022; Irons et al., 2017; Mathôt & Ivanov, 2019; Pilipenko & Samaha, 2024). However, the mechanisms underlying this relationship remain debated: changes in detection rates could reflect genuine gains in perceptual sensitivity or, alternatively, strategic shifts in decision criterion, or changes in subjective signal-likeness (Samaha et al., 2020). By using a multi-alternative forced-choice (m-AFC) partial-report task that is largely criterion-free and in which performance depends on the temporal persistence of sensory information rather than stimulus visibility, our paradigm provides a more direct test of how spontaneous arousal modulates temporal availability of sensory information.

Our results demonstrate that spontaneous fluctuations in pre-stimulus pupil-linked arousal systematically modulate iconic memory performance. Larger pre-stimulus pupil dilation was associated with higher initial stimulus availability (Figure 2A, 3A), leading to improved performance at short stimulus– cue onset asynchronies. At the same time, larger pupils predicted a faster decay of the iconic memory trace (Figure 2A, 3A). In contrast, the asymptotic parameter *a*_0_, reflecting the amount of information transferred into short-term memory, was unaffected by pre-stimulus pupil size. This suggests that arousal specifically influences the transient sensory persistence of visual information, but not the later, more stable short-term memory store. We found no evidence for an inverted U-shaped relationship between arousal and performance, which may reflect the limited arousal range captured in our study. Specifically, spontaneous fluctuations in pupil size likely covered only the lower and intermediate portions of the arousal–performance curve, without reaching the high-arousal “branch” that typically produces performance declines (Aston-Jones & Cohen, 2005). Together, these findings extend previous observations of arousal-related performance benefits in visual detection tasks to the domain of iconic memory, demonstrating that spontaneous arousal modulates the temporal availability of sensory information even in a paradigm largely free from decision-bias confounds.

The effect of arousal on initial stimulus availability mirrors findings from studies using emotional stimuli. Both positively and negatively valenced stimuli have been shown to enhance initial stimulus availability in iconic memory (Kattner & Clausen, 2020; Kuhbandner et al., 2011). This enhancement is thought to reflect the prioritized selection of emotionally salient information from iconic memory. Importantly, emotional stimuli also induce changes in arousal, suggesting that their beneficial effect on early visual persistence may arise, at least in part, from arousal-related mechanisms, consistent with the spontaneous arousal effects observed here.

The faster decay of the memory trace observed with larger pre-stimulus pupil dilation (Figure 2A, 3A) aligns with previous findings on the influence of cortisol on iconic memory (Miller et al., 2015). Elevated cortisol levels, which index heightened arousal, have been shown to accelerate the decay of the iconic trace (Miller et al., 2015). This accelerated decay may reflect an adaptive mechanism linked to the temporal organization of perception. Perception must balance two complementary demands: temporal integration, which promotes perceptual stability by combining visual information over time, and temporal segregation, which enables rapid updating at the cost of stability (Ronconi et al., 2017). This balance is thought to be mediated by oscillatory dynamics that define temporal windows of integration and segregation (Ronconi et al., 2017; Wutz et al., 2014). Arousal may additionally tip this balance toward segregation by speeding up the decay of the iconic trace, thereby promoting updating, reducing interference from earlier stimuli, and enabling rapid processing of new, potentially salient input.

We did not find any effects of pre-stimulus pupil size on confidence ratings (Figure 2B, 3B). This contrasts with previous studies in near-threshold detection tasks, where larger pupil size, indicating higher arousal, has been associated with increased confidence (Pilipenko & Samaha, 2024). The discrepancy likely reflects differences in how confidence is generated across paradigms. In yes/no detection tasks, confidence mainly reflects decision uncertainty, that is, the distance between the internal decision variable and the decision criterion on a given trial, which makes it sensitive to criterion shifts. In contrast, in a multi-alternative forced-choice (m-AFC) task, confidence primarily tracks the strength of the selected item’s representation and therefore covaries with objective accuracy (Peters, 2022). As a result, there is less opportunity for arousal-induced criterion shifts to influence confidence independently of performance. These results open several promising directions for future research. One important step will be to relate the effects of arousal on iconic memory to other neurophysiological measures known to influence sensory persistence. Previous work has shown that strong pre-stimulus alpha power, which is correlated with pupil fluctuations and cortical excitability (Pilipenko & Samaha, 2024; Ruuskanen et al., 2025), enhances initial stimulus availability in iconic memory (Smith & Busch, 2025). The present findings complement this evidence: increased pre-stimulus pupil dilation, reflecting greater arousal, affected the *decay* of iconic memory, whereas pre-stimulus alpha power did not, suggesting two distinct but potentially interacting mechanisms. Future studies could also examine other physiological indicators of arousal, such as respiration, which covaries with both pupil dilation and alpha power and has been shown to modulate visual perception (Kluger et al., 2021, 2024).

Another avenue concerns the interplay between arousal and attention. Arousal has been linked to an increased breadth of attention (Hoeks & Ellenbroek, 1993; Irons et al., 2017; Kolnes et al., 2024), raising the possibility that broader attentional focus contributes to the performance benefits observed in the partial-report paradigm. However, the role of attention in iconic memory remains debated. Some studies suggest that attention modulates iconic memory performance (Botta et al., 2019; Mack et al., 2015, 2016), whereas others argue that iconic memory operates largely independently of attention (Aru & Bachmann, 2017; Bachmann & Aru, 2016; Pinto et al., 2017).

## Conclusion

Our study demonstrates that spontaneous pre-stimulus fluctuations in arousal, indexed by pupil diameter, shape the dynamics of iconic memory. Higher arousal enhances the initial availability of visual information but also accelerates its decay, indicating that arousal modulates the temporal window of sensory persistence. These findings advance our understanding of how moment-to-moment variations in arousal influence both perception and sensory memory.

## Acknowledgments

We thank Teresa Berther, Milena Carolin Koch, Seray-Ezgi Öztekin, Mathilde Maria Pöppelmann, Johanna Seroka, Simon Steibel, and Charlotte Wellmann for help with the data acquisition. We also thank Maximilian Bruchmann for providing us with technical equipment.

